# ChromeCRISPR - A High Efficacy Hybrid Machine Learning Model for CRISPR/Cas On-Target Predictions

**DOI:** 10.1101/2025.04.16.649183

**Authors:** Amirhossein Daneshpajouh, Megan Fowler, Kay C. Wiese

## Abstract

Genome editing has the potential to treat genetic disorders at the source. This can be achieved by modifying the defective DNA through the intentional insertion, deletion, or substitution of genomic content. Among all genome editing technologies, CRISPR/Cas (Clustered Regularly Interspaced Short Palindromic Repeats and CRISPR-associated protein) is considered the gold standard. CRISPR/Cas uses a single guide RNA (sgRNA) to direct the Cas nuclease to a target DNA region. Due to the ease at creating small RNA molecules, it is possible to have the CRISPR/Cas complex target any arbitrary DNA sequence, thus making it a versatile tool. The efficacy of the complex is dependent on the ability of the sgRNA to bind to a complementary DNA sequence, which varies based on the sequence. Thus, a major challenge is finding sgRNA sequences that have good efficacy. This is where computational models can aid scientists: by predicting the activity of sgRNAs to help narrow the search space of finding the optimal sgRNA. We have used a large new dataset to build and compare the ability of several different machine learning architectures’ ability to predict on-target CRISPR/Cas activity. Additionally, we explored how adding GC content affects our sgRNA activity predictions. Our novel hybrid model, ChromeCRISPR, combines the strengths of Convolutional Neural Networks (CNN) with Recurrent Neural Network (RNN) models, has outperformed state-of-the-art models, including DeepHF and AttCrispr, establishing a new benchmark for predictive accuracy in CRISPR/Cas9 efficacy predictions.

## Introduction

CRISPR/Cas has revolutionized genome editing, allowing for precise and versatile modifications to an organism’s DNA. Originally discovered as the adaptive immune system of bacteria and archaea [1, 2], CRISPR/Cas was repurposed into a powerful genome editing tool [3]. It has since been applied to several different applications, ranging from therapeutic gene editing, such as treating *β*-thalassemia and sickle cell disease [4], to agricultural advancements and creating disease-resistant crops [5].

The Clustered Regularly Interspaced Short Palindromic Repeats (CRISPR) and CRISPR-associated proteins (Cas) complex comprises a single guide RNA (sgRNA) and a nuclease (Cas) [6]. The sgRNA consists of two components: a stem-loop structure that binds it to the Cas protein and a guide sequence that directs the complex to a complementary DNA region [6]. When the complex finds a target DNA sequence that is sufficiently complementary, the Cas nuclease induces a double-stranded break [6]. Following this, the homology-directed repair pathway mends the break and incorporates the intended edit [6]. This tool allows researchers to target a specific DNA sequence by creating a complementary sgRNA.

While CRISPR/Cas technology has seen widespread use, predicting the activity of the CRISPR/Cas complex has been challenging. For one, it is important to have high efficacy, that is, the proportion of the number of target sites that have incorporated the modification (on-targets) should be high relative to the number of target sites that were unsuccessfully edited. The efficacy is affected by several factors, including sequence and epigenetic features [7], as well as secondary structure characteristics [8], thus making it difficult to predict the efficacy of a given sgRNA.

Equally, it is vital that the Cas-sgRNA complex has high specificity. There is some flexibility in how complementary the DNA sequence must be to the sgRNA in order to induce a modification, so it is possible for non-target DNA regions to be edited. These off-target modifications pose a risk to the safety and reliability of CRISPR/Cas genome editing.

Thus, accurate prediction of CRISPR/Cas activity is an essential step in ensuring the continued development of the CRISPR/Cas technology. Assessing the efficacy and specificity of sgRNA activity experimentally is time-consuming and resource-intensive, thus many have turned to computational approaches in order to prioritize biological validation of only the most promising sgRNAs.

Several computational approaches exist for this task, although these tools have their own set of challenges. As mentioned, running CRISPR/Cas activity assays in any great quantity is intractable and due to this, there is little data available to train computational models. This results in models with limited predictive power and restricted size and complexity.

Additionally, the development of computational models for predicting off-target sites is approached as a separate endeavor from on-target predictive models, driven by the necessity to harness distinct datasets. In our recent paper, we explore off-target models, producing novel insights and methodologies [9]. Particularly, our research looks at the application of evolutionary algorithms in the domain of off-target prediction.

The work presented here expands upon our previous work in which we assessed the performance of five baseline models: A Random Forest, a CNN (Convolutional Neural Network), and three RNNs (Recurrent Neural Networks) [10]. For one of these RNNs, we also demonstrated how adding GC content could be worthwhile for improving our predictions. Here, we review these baseline models, and then we build upon them to show the effect of adding GC content to the other models, as well as evaluating how the depth of the models affects the performance. We then combine these techniques to create CNN-RNN hybrid models with GC content, which we have collectively named ChromeCRISPR (**C**NN **H**ybridized **R**NN **O**n-target **M**odels for Gene **E**diting with **CRISPR**). Herein, we show that our best ChromeCRISPR model, a CNN-GRU (Gated Recurrent Unit) hybrid model, outperforms other current state-of-the-art techniques developed from the same dataset. Of these other techniques, DeepHF [11] had previously achieved a Spearman correlation of 0.867 and mean squared error of 0.0094, while AttCRISPR [12] attained a Spearman correlation of 0.872 (mean squared error not reported). Our ChromeCRISPR model establishes new benchmarks for both Spearman correlation and mean squared error on this dataset, recording 0.876 and 0.0093 respectively.

The significance of this study lies in its potential to contribute to the future success of CRISPR/Cas-based therapies for genetic disorders, as well as gaining insights into which machine learning techniques are best suited for this task. Designing highly potent sgRNAs that specifically target the defective DNA will lead to more precise and safe gene editing, thereby facilitating the development of treatments for previously incurable genetic diseases. From the computational perspective, we show the power of hybrid models, as well as the impact of adding GC content as a feature. The establishment of robust predictive models for CRISPR/Cas activity promises a transformative impact on human health and medical advancements.

## 1 Literature Review

Several tools exist for predicting on-target activity for CRISPR/Cas genome editing. Over the last decade, these tools have seen a great deal of development, beginning as empirical rule-based approaches based on biological experiments [7, 13–16]. These initial tools focused on nucleotide positions and a handful of thermodynamic properties including GC content and the melting temperature.

With the rise in popularity of machine learning, there was a surge in traditional machine learning methods such as regressive and support vector machine (SVM) based methods [14, 17–19]. This was closely followed by more contemporary methods using deep learning and neural networks [11, 20–27]. The deep learning and neural network models were mostly convolutional neural networks (CNNs), however, DeepHF [11] opted to use a recurrent neural network (RNN) to good effect. A detailed account of these tools is given in a recent review by Sherkatghanad et al. [28].

One of the main challenges faced by all these models is the overall lack of data availability. One of the reasons that it is important to develop predictive models is that it is time-consuming and resource-intensive to test many sgRNAs for their activity, however, this has also resulted in a deficiency in the amount of data available for training large models. Combining datasets across studies is also often unfeasible, since collecting CRISPR datasets involves the compilation of data from diverse cell types and organisms, assessed through various biological assays. This amalgamation results in a heterogeneous dataset and may introduce batch effects, where non-biological factors in an experiment influence the data. Thus to date, models have been limited in their size.

Fortunately, one of the recent on-target predictive tools also conducted a wide-scale experiment for sgRNA on-target activity [11]. This study evaluated the activity of nearly 60,000 sgRNAs across approximately 20,000 genes. This dataset has been used by a few different models, including CRISPRpred(SEQ) [19], CRISPR-ONT [25], AttCRISPR [12], and the group who developed the dataset’s own tool, DeepHF [11]. However, only DeepHF [11] and AttCRISPR [12] trained and evaluated their models exclusively with this dataset.

In the continuous evolution of CRISPR/Cas9 efficacy prediction, the introduction of AttCRISPR [12] marks a notable advancement, by integrating attention mechanisms, a concept borrowed from the field of natural language processing. It leverages a temporal attention module that dynamically assigns significance to different segments of the sgRNA sequence, enhancing the model’s focus on critical nucleotides that influence on-target activity. This mechanism operates through a sophisticated interplay of queries, keys, and values also using bidirectional Gated Recurrent Unit (GRU) networks, enabling a nuanced understanding of sequence determinants. Furthermore, AttCRISPR adopts an ensemble approach, incorporating biologically relevant features such as secondary structure and GC content, processed alongside direct sequence inputs. By combining these elements through a stacking strategy, AttCRISPR achieves a balance of high predictive accuracy and interpretability, making it a valuable tool for sgRNA design and optimization in the CRISPR/Cas9 system.

We sought to build upon these previous works and began exploring the predictive ability of several different machine learning methods in [10]. There, we assessed a Random Forest, a CNN (Convolutional Neural network), a GRU (Gated Recurrent Network), an LSTM (Long Short-Term Memory), and a BiLSTM (Bi-directional LSTM). We discovered that the Random Forest performed significantly worse than the other models, and that the three RNNs significantly outperformed the CNN. We then added GC content as a feature to our LSTM to determine if this may improve its performance at predicting the activity of sgRNAs on either end of the GC content spectrum. These results proved promising, and thus we have expanded upon the idea in this paper.

Here, we evaluate the use of GC content for more of our models, and we also assess the effect of deeper models. Inspired by the success of hybrid CNN-RNN models like C-RNNCrispr [24], we experimented with the hybrid models in our work as well. We combined the CNN model with its powerful feature extraction capabilities with an RNN to achieve a high performing model, ChromeCRISPR, that outperformed the top models from both DeepHF [11] and AttCRISPR [12].

## 2 Methods

We employ several different analytical, processing, and predictive technologies. In the following sections we describe our method for analyzing the DeepHF raw data, our processing steps, which include adding features and encoding the biological data into a numerical format, and our general approach for hyperparameter tuning and training. We then discuss the various models that we evaluated followed by the evaluation techniques used.

### 2.1 Data Analysis

The data utilized in this study were sourced from the DeepHF study [11]. This dataset contains activity values for almost 60,000 unique sgRNAs compiled from 20,000 human genes. The study compiled the CRISPR/Cas activity values for three different Cas enzymes: the wildtype SpCas9, eSpCas9, and SpCas9-HF. However, we have elected to only use the wildtype SpCas9 data for our model since it is the standard Cas enzyme and the one most commonly used. Additionally, we elected not to combine the wildtype SpCas9 data with the eSpCas9 and SpCas9-HF data due to the fact that they have statistically significantly different underlying activity distributions, with the mean values being 0.72, 0.35, and 0.48, respectively. Each sgRNA sequence is 20 nucleotides long plus the variable PAM nucleotide, for a total sequence length of 21 nucleotides.

The activity values are measured as indel (insertion or deletion) frequencies, showing the effectiveness of CRISPR/Cas9 editing. For any given sgRNA (*sgRNA*_*x*_) and its corresponding target DNA sequence (*DNA*_*x*_), the activity value shows the fraction of *DNA*_*x*_ target sites that were successfully edited—shown by insertions or deletions—compared to the total *DNA*_*x*_ target sites looked at. To find this ratio, we divide the number of target DNA molecules that have indels, showing successful gene editing, by the total number of target DNA molecules checked, which includes both edited and unedited sequences:

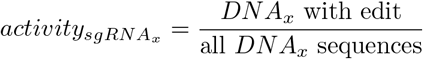

### 2.2 Features

Several biological features have been shown to have an impact on the activity of CRISPR/Cas genome editing including the melting temperatures, free energy, and DNA accessibility [8,29,30]. For this paper we evaluate only the effect of GC content, however as we discuss later, we plan to assess in detail the effects of these other biological features as well in future work.

#### 2.2.1 GC Content

GC content can approximate the thermodynamic stability of an oligonucleotide duplex. Compared to AT/U pairs, GC pairs have an additional hydrogen bond which adds to the stability of the duplex. For CRISPR/Cas experiments, ideal GC content is in the range of 40-60%, where sgRNAs with GC content outside of that range tend to have lower efficiency [31]. This trend of sgRNAs with extreme GC content having low activity was also noted by Doench et al [13] and Wang et al [32]. The GC content for a given sgRNA (*sgRNA*_*x*_) is calculated as the percentage of guanine and cytosine nucleotides in the sequence:

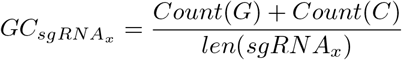

GC content ranged from 10-95% with a mean of 52% and standard deviation of 11%, thus the majority of our sgRNAs had GC content within the 40-60% range indicating higher activity. Within our dataset, we also found that GC content within the 40-60% range was associated with a higher activity (t-test *p <* 0.05).

### 2.3 Encoding

One-hot encoding was used to encode the sgRNA sequences into a numerical representation, as it is a common way to translate DNA and RNA sequences into a numerical representation (Figure 1). Each nucleotide in our 21mer sequence was converted to an array of length 4, thus producing a matrix with dimensions 21x4, where each row is the nucleotide identity and each column represents the position in the sequence. A cell is assigned a value of 1 if at that column position, it contains the row nucleotide, 0 otherwise. This matrix was then flattened to a 1-dimensional array of length 84.

**Figure 1:**
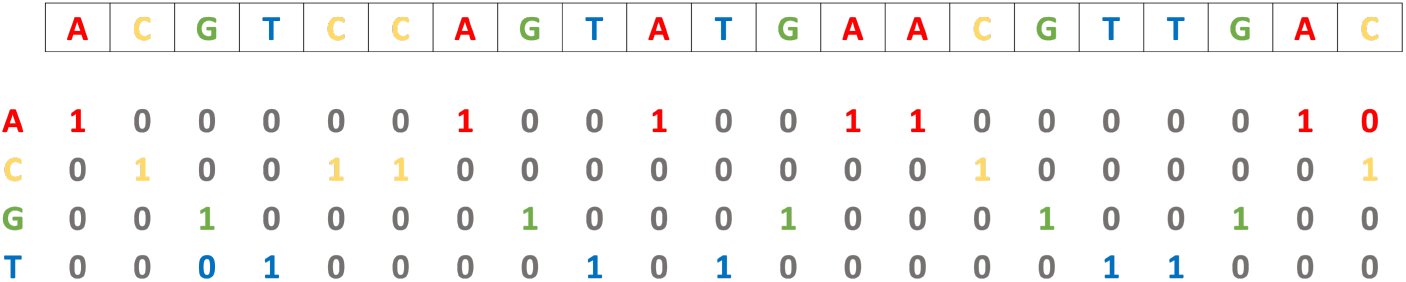
One hot encoding of sgRNA 21mer sequences

### 2.4 Hyperparameter Tuning and Training

We set aside 15% of the data for testing and used the remaining 85% for hyperparameter tuning and training. We used a nested 5-fold cross-validation with Bayesian search for hyperparameter tuning. That is, the 85% of data originally split off was further divided such that we performed five Bayesian searches for the best hyper-parameters on 68% of the data and used the remaining 17% for the validation set. The selected hyperparameters were then validated using 5-fold cross-validation. The model was then retrained on the full 85% portion of the data.

Our models were trained on hardware with 4GB RAM memory and utilized 2 CPU cores. Additionally, we utilized NVIDIA V100 Volta GPUs with 32GB HBM2 memory, deployed on the Digital Research Alliance of Canada superclusters. The training process took approximately 20 seconds per iteration on average.

### 2.5 Models

In this study, we evaluated several machine learning models to predict CRISPR/Cas on-target efficacy. Each model brings a unique approach to handling sequence data, providing insights into their effectiveness for this task. The Random Forest was implemented in scikit learn [33], while the other neural neworks were implemented in PyTorch [34]. A summary of the models we evaluated is given in Table 1.

**Table 1:** Summary of Models.

| Model Group | Model Name |
| --- | --- |
| Base Model | RF |
|  | CNN |
|  | GRU |
|  | LSTM |
|  | BiLSTM |
| Base Model + GC | CNN+GC |
|  | GRU+GC |
|  | LSTM+GC |
|  | BiLSTM+GC |
| Deep Models | deepCNN |
|  | deepGRU |
|  | deepLSTM |
|  | deepBiLSTM |
| Deep Models + GC | deepCNN+GC |
|  | deepGRU+GC |
|  | deepLSTM+GC |
|  | deepBiLSTM+GC |
| ChromeCRISPR (Hybrid CNN Models + GC) | CNN_GRU+GC |
|  | CNN_LSTM+GC |
|  | CNN_BiLSTM+GC |

#### 2.5.1 Random Forest (RF)

The Random Forest model is an ensemble learning method, utilizing multiple decision trees to make predictions. Its strength lies in its ability to reduce overfitting by averaging the results of individual trees. This model is particularly effective in handling non-linear data and provides a good benchmark for more complex models.

#### 2.5.2 Convolutional Neural Network (CNN)

Convolutional Neural Networks, widely used in image recognition tasks, have shown promise in sequence analysis due to their ability to detect patterns and motifs in sequential data. In our CNN model, convolutional layers are used to identify and learn sequence features critical for predicting CRISPR/Cas efficacy.

#### 2.5.3 Gated Recurrent Unit model (GRU)

GRU is a type of recurrent neural network that excels in learning from sequential data, such as nucleotide sequences. It is particularly efficient due to its simplified architecture, which helps in faster training without significant compromise in performance, especially in smaller datasets.

#### 2.5.4 Long Short-Term Memory Network (LSTM)

LSTM, another variant of recurrent neural networks, is known for its ability to capture long-term dependencies in sequence data, an important factor considering the dependencies in nucleotide sequences for CRISPR/Cas targeting efficacy. It addresses the issue of vanishing gradients which is common in traditional RNNs.

#### 2.5.5 Bidirectional Long Short-Term Memory Network (BiLSTM)

BiLSTM extends the capabilities of standard LSTM by processing the data in both forward and reverse directions. This approach is beneficial for capturing context from both ends of the sequence, potentially improving the model’s ability to learn complex patterns in sgRNA sequences.

#### 2.5.6 Transformer

Transformer, known for their effectiveness in natural language processing tasks, employ self-attention mechanisms to weigh the significance of different parts of the input data. In the context of sgRNA sequences, this model can help in identifying key nucleotides and motifs that are crucial for targeting efficacy.

### 2.6 Evaluation Metrics

To assess the performance of the models, we employed two key metrics: Mean Squared Error (MSE) and Spearman Correlation Coefficient. Scikit-learn’s mean squared error [33] was used to calculate the MSE and SciPy [35] was used to calculate the Spearman correlation.

#### 2.6.1 Mean Squared Error (MSE)

MSE is a widely used metric for regression models. It measures the average of the squares of the errors, i.e., the average squared difference between the estimated values and the actual value. A lower MSE value indicates a better fit of the model to the data. In the context of our study, it quantifies the difference between the predicted efficacy of sgRNAs and their actual efficacy observed in experimental data.

#### 2.6.2 Spearman Correlation Coefficient

The Spearman correlation is a non-parametric measure of rank correlation. It assesses how well the relationship between two variables can be described using a monotonic function. In our study, this metric is crucial for understanding the strength and direction of the association between the predicted and observed sgRNA efficacies. A higher Spearman correlation indicates that the model predictions are more closely aligned with the actual efficacy ranks of sgRNAs.

These metrics provide a comprehensive view of the models’ performance, considering both the accuracy of predictions (MSE) and the rank-order consistency (Spearman Correlation) with the observed data.

### 2.7 Testing

Our test set was composed of 15% of the data and included predictions for 8341 sgRNAs. When testing each model, we performed 10-fold cross validation.

### 2.8 Statistical Analysis

One-way ANOVA [36] was used to test for significant differences between the means of various groups, such as between models. Following a significant result, Tukey’s Honest Significant Difference (HSD) [37] was used to extract which means were significantly different than the others of that group.

## 3. Results

### 3.1 Comparison of Machine Learning Models

We began by testing out various machine learning model architectures to ascertain if one type performed better than the others (Figure 2). We tested a Random Forest, a Convolutional Neural Network (CNN), and three different implementations of a Recurrent Neural Network (RNN): A Gated Recurrent Unit (GRU) model, a Long Short-Term Memory (LSTM) model, and a Bidirectional Long Short-Term Memory (BiLSTM) model. We found that in general, the RNNs performed the best, having both the lowest mean squared error (MSE) (Figure 2a) and highest Spearman correlation (Figure 2b).

**Figure 2:**
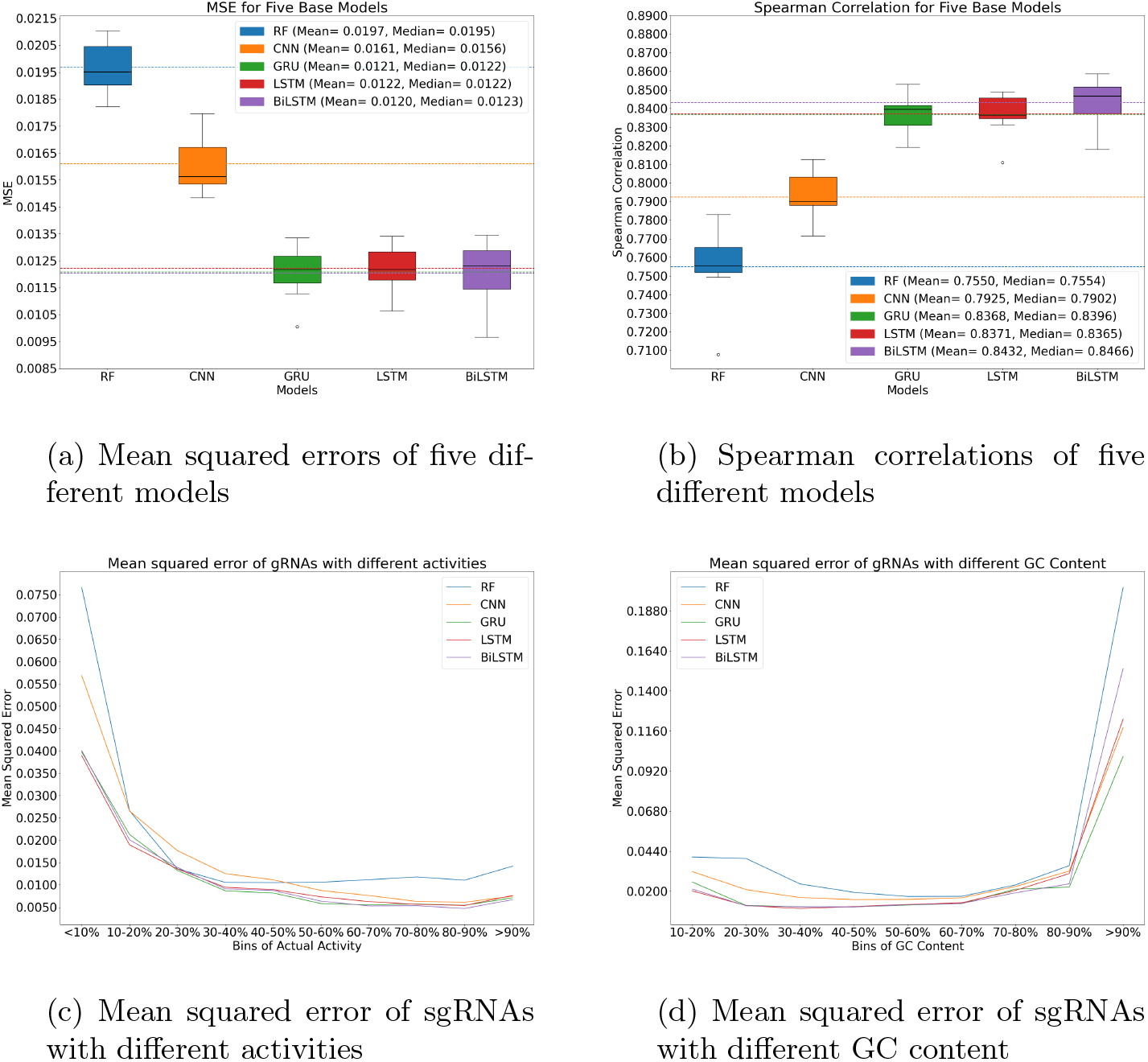
Comparison of five different base models. **a** The mean squared error of the RF, CNN, GRU, LSTM, and BiLSTM models. The horizontal dashed lines represent the means, while the black solid line in each box plot is the median. The mean MSEs and median MSEs for the models are RF = 0.0197 and 0.0195, CNN = 0.0161 and 0.0156, GRU = 0.0121 and 0.0122, LSTM = 0.0122 and 0.0122, and BiLSTM = 0.0120 and 0.0123, respectively. **b** The spearman correlation of the RF, CNN, GRU, LSTM, and BiLSTM models. The horizontal dashed lines represent the means, while the black solid line in each box plot is the median. The mean spearman correlations and median spearman correlations for the models are RF = 0.7550 and 0.7554, CNN = 0.7925 and 0.7902, GRU = 0.8368 and 0.8396, LSTM = 0.8371 and 0.8365, and BiLSTM = 0.8432 and 0.8466, respectively. **c** The mean squared error of the RF, CNN, GRU, LSTM, and BiLSTM across different deciles of sgRNA activity. **d** The mean squared error of the RF, CNN, GRU, LSTM, and BiLSTM across sgRNAs with varying GC content.

As a posthoc analysis, we then sought to inspect these results further to try to elucidate where our models performed well and where they struggled. First, due to the dataset skewing towards higher activity values, we thought that our model may perform better on sgRNAs in this range. We binned the test set by their true activity deciles, that is, the sgRNAs in the bottom 10% for activity were binned into the *<* 10% activity bin, and so forth. Our hypothesis proved correct, in that our model struggled to predict the activity of sgRNAs in the bottom 30% (Figure 2c). We next examined how our model fared on sequences of different GC content. We again binned the sgRNA test set by GC content, however, this time, we did so by the value of the GC content. That is, if a sgRNA had a GC content between 0-10% it was binned in the *<* 10% GC bin. In general, we found the predictions to be consistent across GC content, except for the very high (*>* 90%) GC content sgRNAs (Figure 2d). We thought that this could be due to sgRNAs with high GC content having low activity, as then we could attribute this phenomenon to the data imbalance. However, we found that in our dataset, sgRNAs with high GC content were actually associated with high activity (Spearman = 0.3062, *p <* 0.05). Thus we can address these two areas of poor predictive performance separately.

### 3.2 Adding GC Content as a Feature

We decided to focus on improving the predictive performance of our models by addressing the poor performance on sgRNAs with high GC content. To this end, we added the GC content as a feature into our neural network models. We obtained mixed results for this experiment, as the CNN and GRU showed a decrease in performance, with a higher MSE (CNN: 0.0161 to 0.0170, GRU: 0.0121 to 0.0122) and lower Spearman (CNN: 0.7925 to 0.7810, GRU: 0.8368 to 0.8401), however we found that neither of these differences were statistically significant (*p >* 0.05) (Tables 2 & 3). Conversely, the LSTM and BiLSTM both had an increase in performance, with a lower MSE (LSTM: 0.0122 to 0.0112, BiLSTM: 0.0120 to 0.0110), but again, this change was not statistically significant (*p >* 0.05) (Tables 2 & 3).

**Table 2:** MSE comparison across all models.

| Model Name | Base Model | Model+GC | deepModel | deepModel+GC | *Hybrid+GC |
| --- | --- | --- | --- | --- | --- |
| CNN | 0.0161 | 0.0170 | 0.0098 | 0.0093 | — |
| GRU | 0.0121 | 0.0122 | 0.0099 | 0.0098 | **0.0093** |
| LSTM | 0.0122 | 0.0112 | 0.0103 | 0.0104 | 0.0115 |
| BiLSTM | 0.0120 | 0.0110 | 0.0104 | 0.0098 | 0.0096 |

\*Hybrid models are CNN followed by the RNN model. See Table 1.

**Table 3:** Spearman comparison across all models.

| Model Name | Base Model | Model+GC | deepModel | deepModel+GC | *Hybrid+GC |
| --- | --- | --- | --- | --- | --- |
| CNN | 0.7925 | 0.7810 | 0.8694 | 0.8728 | – |
| GRU | 0.8368 | 0.8401 | 0.8684 | 0.8668 | **0.8760** |
| LSTM | 0.8371 | 0.8564 | 0.8620 | 0.8602 | 0.8668 |
| BiLSTM | 0.8432 | 0.8550 | 0.8617 | 0.8671 | 0.8700 |

\*Hybrid models are CNN followed by the RNN model. See Table 1.

Comparing our models to one another, we found a similar pattern as was observed previously, with the RNNs significantly outperforming the CNN in the MSE (*p <* 0.05) as well as having higher Spearman correlations (Figures 3a & 3b). Of note, across the three RNN models, less than 0.5% of the predictions were off by more than 5%, in contrast with the CNN which was off by more than 5% for 0.74% of the predictions.

**Figure 3:**
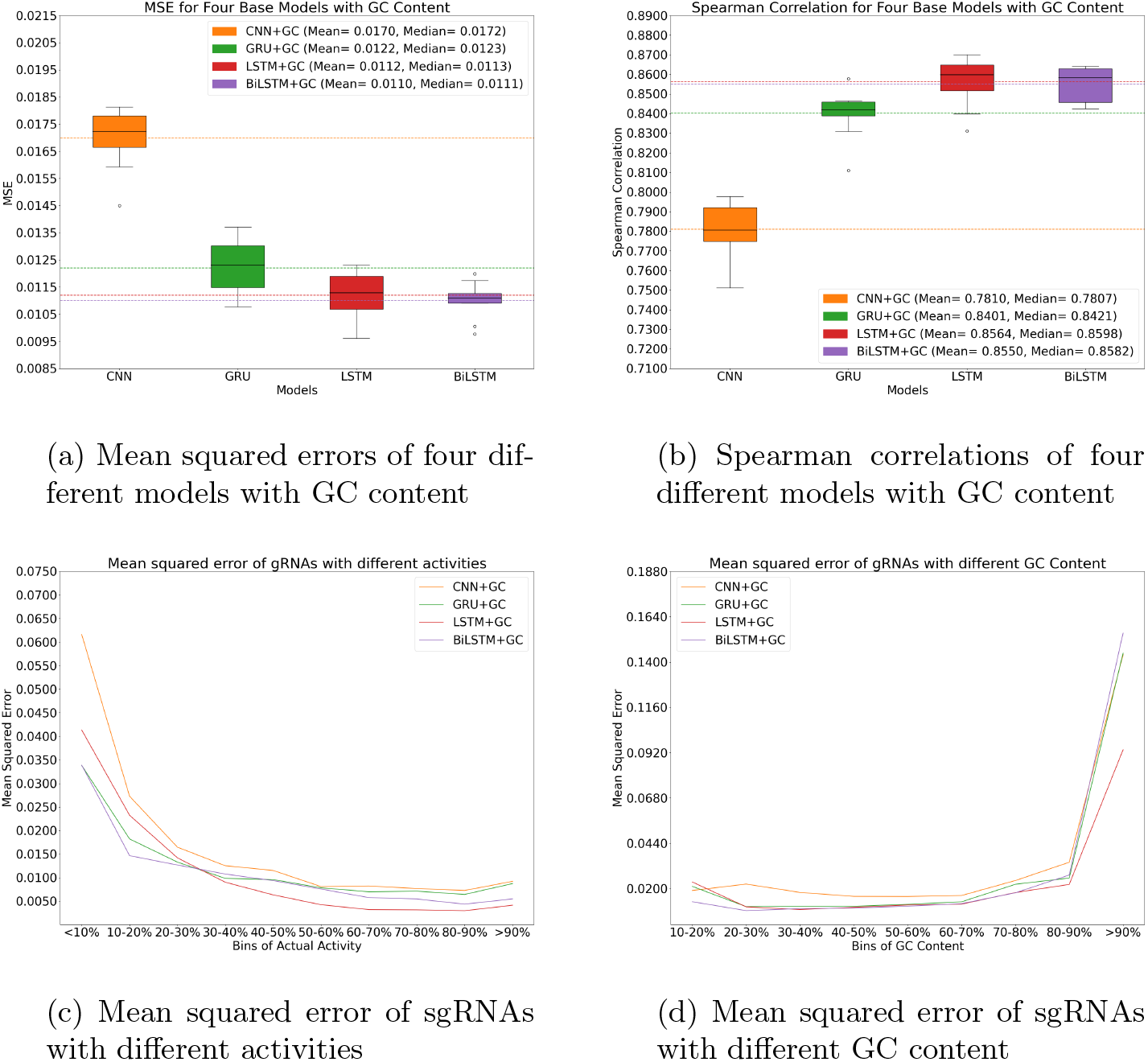
Comparison of four different base models with GC content added as a feature. **a** The mean squared error of the CNN+GC, GRU+GC, LSTM+GC, and BiLSTM+GC models. The horizontal dashed lines represent the means, while the black solid line in each box plot is the median. The mean MSEs and median MSEs for the models are CNN+GC = 0.0170 and 0.0172, GRU+GC = 0.0122 and 0.0123, LSTM+GC = 0.0112 and 0.0113, and BiLSTM+GC = 0.0110 and 0.0111, respectively. **b** The Spearman correlation of the CNN+GC, GRU+GC, LSTM+GC, and BiLSTM+GC models. The horizontal dashed lines represent the means, while the black solid line in each box plot is the median. The mean Spearman correlations and median Spearman correlations for the models are CNN+GC = 0.7810 and 0.7807, GRU+GC = 0.8401 and 0.8421, LSTM+GC = 0.8564 and 0.8598, and BiLSTM+GC = 0.8550 and 0.8582, respectively. **c** The mean squared error of the CNN+GC, GRU+GC, LSTM+GC, and BiL-STM+GC across different deciles of sgRNA activity. **d** The mean squared error of the CNN+GC, GRU+GC, LSTM+GC, and BiLSTM+GC across sgRNAs with varying GC content.

Similarly to our post hoc analysis of the base models, we binned the sgRNAs by their true activity and by GC content (Figures 3c & 3d). In this case, the BiLSTM showed the best performance on the sgRNAs with lower activity (0−30%), while the LSTM had the best performance for both the mid range and highly active sgRNAs (Figure 3c). Interestingly, despite performance improving on the low activity, we did not see any statistically significant improvement across any of the GC bins (Figure 3d).

### 3.3 Deeper Models

Adding more layers to a neural network architecture does not necessarily improve performance beyond a certain point. This phenomenon, often referred to as over-fitting, occurs when the model becomes too complex and starts to fit the noise in the training data rather than the underlying patterns. In the case of our experiments, we observe that beyond a certain number of layers, both the test loss and the test Spearman score plateau or even degrade, indicating that the additional complexity does not lead to better generalization performance on unseen data (Figure 4). Therefore, it is crucial to carefully balance model complexity with performance metrics to avoid overfitting and ensure optimal model performance.

**Figure 4:**
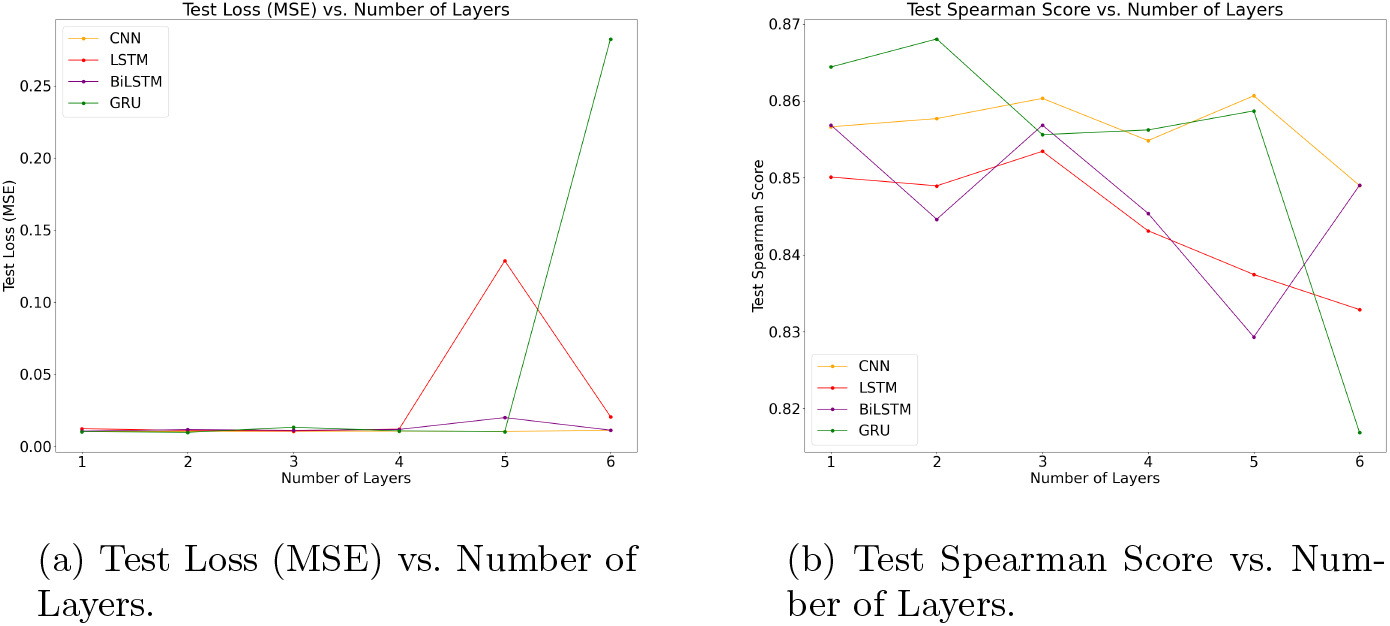
Comparison of test loss (MSE) and test Spearman score vs. number of layers in neural network architectures. The left plot (a) shows how the test loss (MSE) varies with the number of layers. The right plot (b) illustrates the relationship between the test Spearman score and the number of layers. In both plots, it is observed that increasing the number of layers initially improves performance, but beyond a certain point, the performance stabilizes or deteriorates, indicating diminishing returns or potential overfitting.

We then added the number of layers as a hyperparameter and retrained the models. Following this, we tested our models again and found that both the deepCNN and deepGRU had a statistically significant improvement in performance over both the base model and the base model with GC content added (*p <* 0.05). We also found statistically significant improvement of the deepLSTM over the base LSTM and of the deepBiLSTM over the base BiLSTM (*p <* 0.05).

These deeper models all performed similarly, including the CNN which saw a drastic improvement in performance from the base models (Figures 5a & 5b). The GRU also had a slight edge at predicting sgRNAs in the lower range of activities, though this was not significant (*p >* 0.05), but at higher sgRNA activities, the LSTM and CNN significantly outperformed the GRU (Figure 5c).

**Figure 5:**
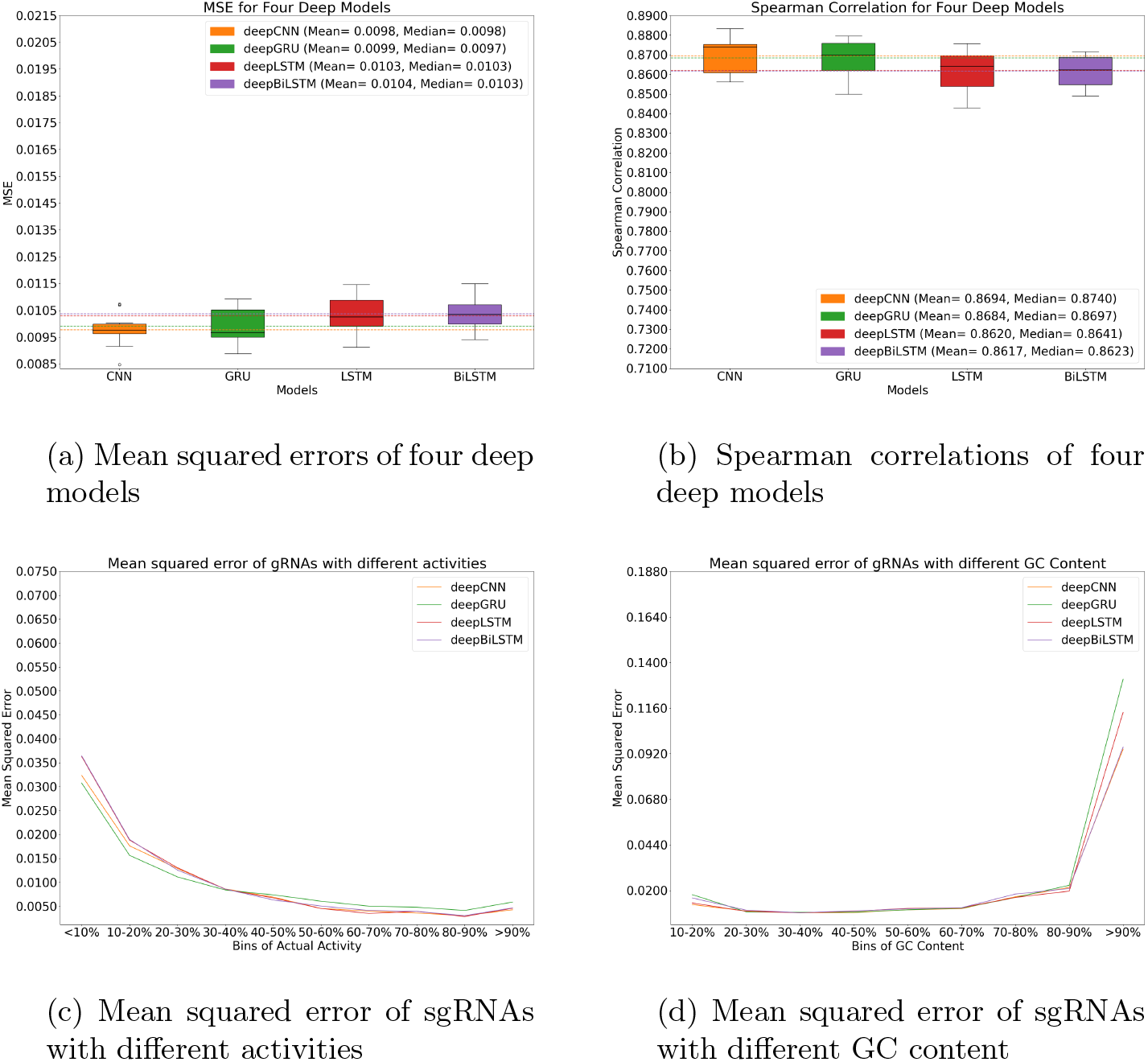
Comparison of four deep Models. **a** The mean squared error of the deep-CNN, deepGRU, deepLSTM, and deepBiLSTM models. The horizontal dashed lines represent the means, while the black solid line in each box plot is the median. The mean MSEs and median MSEs for the models are deepCNN = 0.0098 and 0.0098, deepGRU = 0.0099 and 0.0097, deepLSTM = 0.0103 and 0.0103, and deepBiLSTM = 0.0104 and 0.0103, respectively. **b** The Spearman correlation of the deepCNN, deepGRU, deepLSTM, and deepBiLSTM models. The horizontal dashed lines represent the means, while the black solid line in each box plot is the median. The mean Spearman correlations and median Spearman correlations for the models are deepCNN = 0.8694 and 0.8740, deepGRU = 0.8684 and 0.8697, deepLSTM = 0.8620 and 0.8641, and deepBiLSTM = 0.8617 and 0.8623, respectively. **c** The mean squared error of the deepCNN, deepGRU, deepLSTM, and deepBiLSTM across different deciles of sgRNA activity. **d** The mean squared error of the deepCNN, deepGRU, deepLSTM, and deepBiLSTM across sgRNAs with varying GC content.

Interestingly, the CNN performed statistically significantly better than the base model and base model with GC content on the sgRNAs in the lowest 100 for GC content (*p <* 0.05). This trend was not repeated for the RNN models, which did not show any significant changes in the sgRNAs with lower GC content or higher GC content.

### 3.4 Deeper Models with GC Content as a Feature

Since we did see some improvement to the base models using GC content, we opted to assess how this would affect our deeper models. Again, we detected no significant difference in the MSE predictions of any of the four models (Figures 6a & 6b). We also were unable to detect any significant difference in the way each model predicts the activity deciles of the sgRNAs (Figure 6c) and the models all showed similar performance across sgRNAs with differing amounts of GC content (Figure 6d).

**Figure 6:**
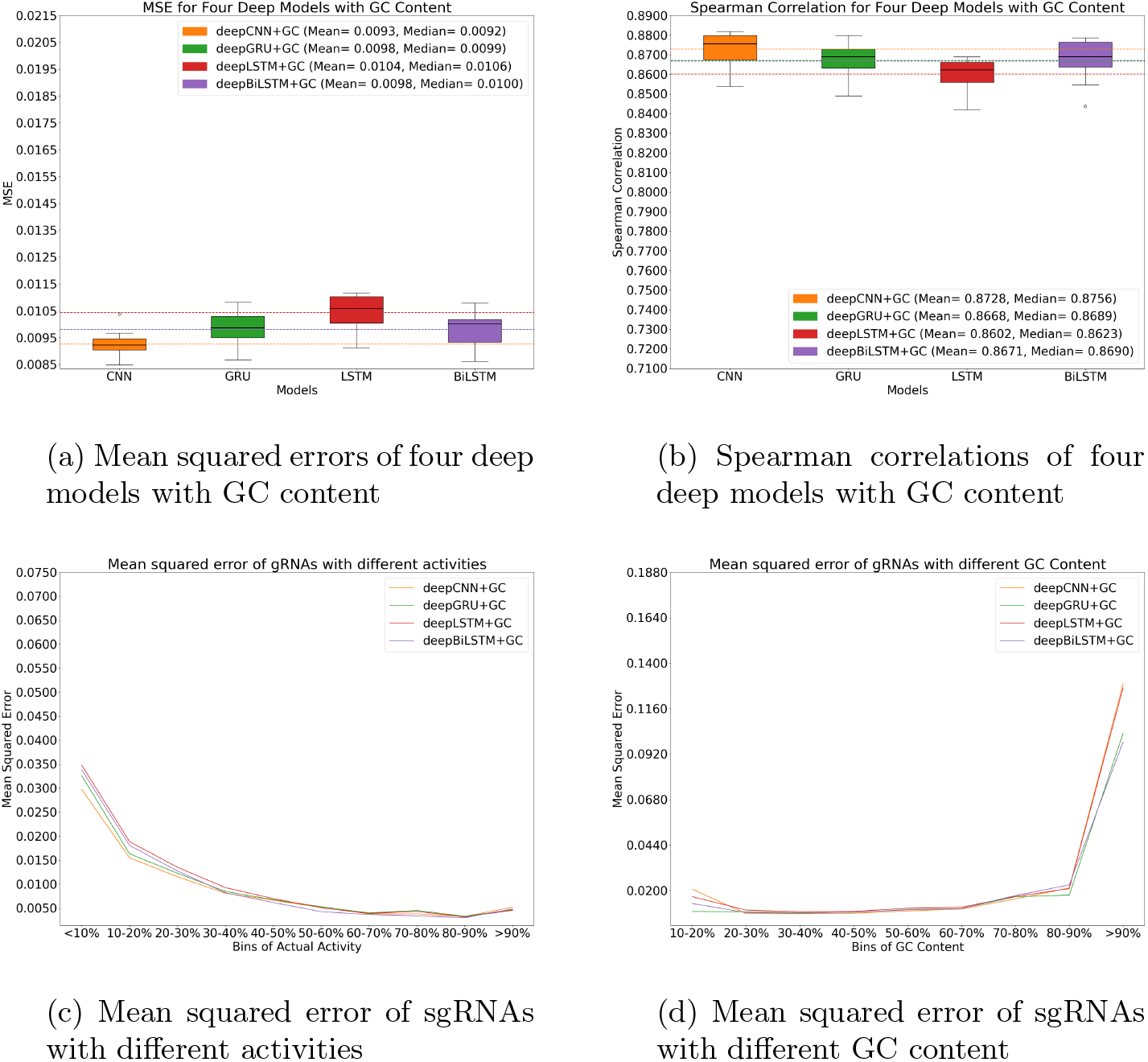
Comparison of four deep models with GC Content added as a feature. **a** The mean squared error of the deepCNN+GC, deepGRU+GC, deepLSTM+GC, and deepBiLSTM+GC models. The horizontal dashed lines represent the means, while the black solid line in each box plot is the median. The mean MSEs and median MSEs for the models are deepCNN+GC = 0.0093 and 0.0092, deep-GRU+GC = 0.0098 and 0.0099, deepLSTM+GC = 0.0104 and 0.0106, and deep-BiLSTM+GC = 0.0098 and 0.0100, respectively. **b** The Spearman correlation of the deepCNN+GC, deepGRU+GC, deepLSTM+GC, and deepBiLSTM+GC models. The horizontal dashed lines represent the means, while the black solid line in each box plot is the median. The mean Spearman correlations and median Spearman correlations for the models are deepCNN+GC = 0.8728 and 0.8756, deepGRU+GC = 0.8668 and 0.8689, deepLSTM+GC = 0.8602 and 0.8623, and deepBiLSTM+GC = 0.8671 and 0.8690, respectively. **c** The mean squared error of the deepCNN+GC, deepGRU+GC, deepLSTM+GC, and deepBiLSTM+GC across different deciles of sgRNA activity. **d** The mean squared error of the deepCNN+GC, deepGRU+GC, deepLSTM+GC, and deepBiLSTM+GC across sgRNAs with varying GC content.

### 3.5 ChromeCRISPR Hybrid Models

Since the CNN was now performing similarly to the RNN models, we thought that combining the CNN with an RNN could allow our model to harness the feature extraction capabilities of the CNN while maintaining the sequence processing abilities of the RNN. We therefore tested hybrid models of a CNN linked to each of the three RNNs (Figure 7). We opted to include GC content as a feature once more since despite the RNNs not improving, or even in some cases such as the GRU and LSTM to have reduced performance, the deep CNN did see a slight improvement (Tables 2 & 3).

**Figure 7:**
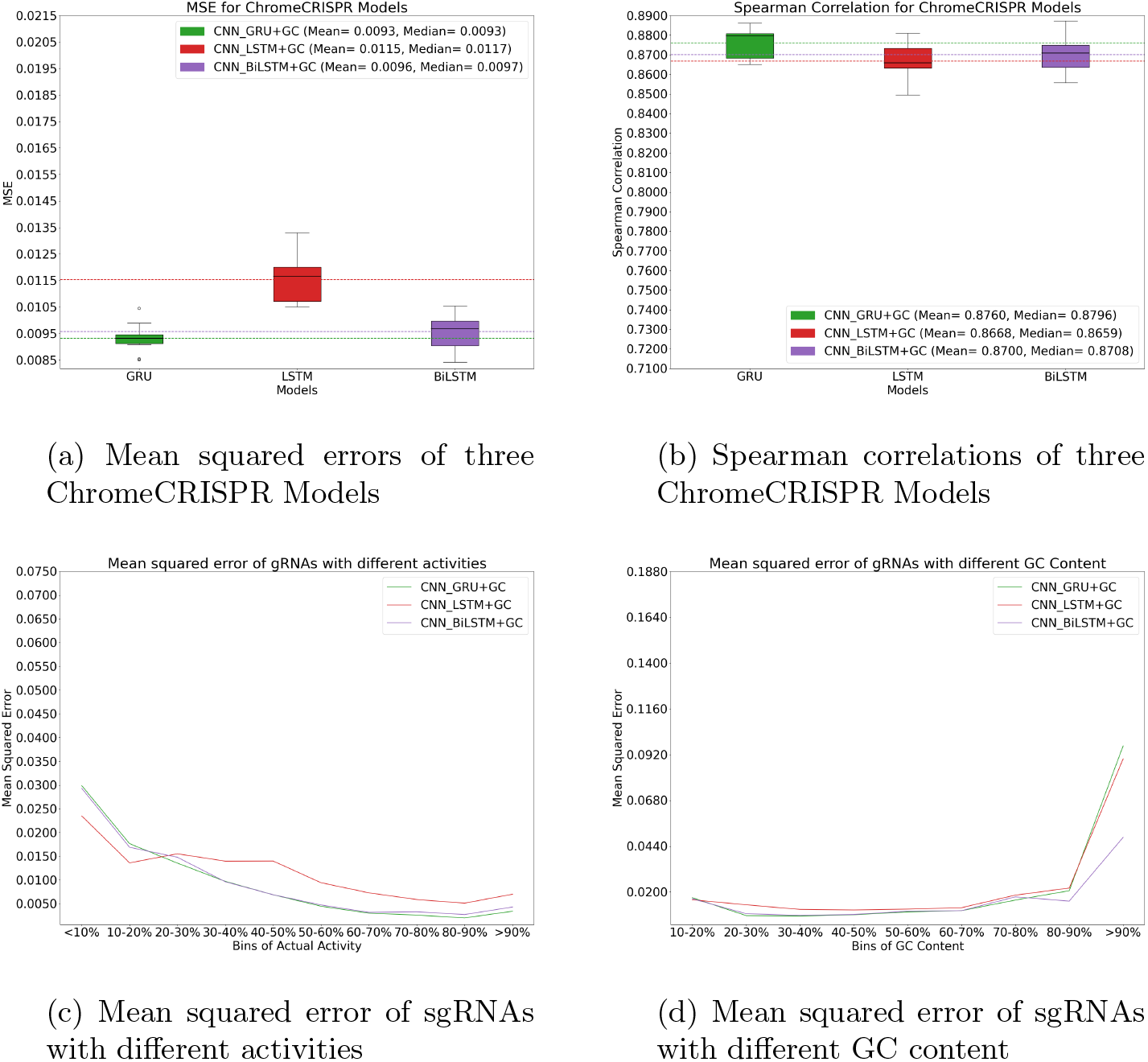
Comparison of three ChromeCRISPR models, each of which is a CNN hybridized RNN model with GC content. **a** The mean squared error of the CNN GRU+GC, CNN LSTM+GC, and CNN BiLSTM+GC models. The horizontal dashed lines represent the means, while the black solid line in each box plot is the median. The mean MSEs and median MSEs for the models are CNN GRU+GC = 0.0093 and 0.0093, CNN LSTM+GC = 0.0115 and 0.0117, and CNN BiLSTM+GC = 0.0096 and 0.0097, respectively. **b** The Spearman correlation of the CNN GRU+GC, CNN LSTM+GC, and CNN BiLSTM+GC models. The horizontal dashed lines represent the means, while the black solid line in each box plot is the median. The mean Spearman correlations and median Spearman correlations for the models are CNN GRU+GC = 0.8760 and 0.8796, CNN LSTM+GC = 0.8668 and 0.8659, and CNN BiLSTM+GC = 0.8700 and 0.8708, respectively. **c** The mean squared error of the CNN GRU+GC, CNN LSTM+GC, and CNN BiLSTM+GC across different deciles of sgRNA activity. **d** The mean squared error of the CNN GRU+GC, CNN LSTM+GC, and CNN BiLSTM+GC across sgRNAs with varying GC content.

The CNN GRU+GC and the CNN BiLSTM+GC models both showed a slight improvement in MSE and Spearman correlation, although the performance of the CNN LSTM+GC dipped. None of these performance deviations from their deep model counterparts were found to be statistically significant (*p >* 0.05). There was however, a statistically significant difference between the performance of the CNN GRU+GC and the CNN BiLSTM+GC models with the CNN LSTM+GC model (Figures 7a & 7b).

We believe a possible explanation for the under-performance of the CNN LSTM+GC model is due to the direction of the sequence in our dataset. As mentioned in section 2.1, our sgRNA sequence is composed of the 20 nucleotide sequence and the variable PAM nucleotide. This sequence is given in the canonical 5’ to 3’ direction, and thus the variable PAM nucleotide is the last nucleotide of the sequence. We therefore hypothesize that the reason the CNN BiLSTM+GC outperforms the CNN LSTM+GC is due to the BiLSTM having access to the PAM nucleotide from the start, whereas the LSTM must traverse the entire sequence before it sees the PAM nucleotide. We believe there could be a similar reason for the CNN GRU+GC’s out-performance. Since the GRU is less able to retain long-term dependencies, when it arrives at the end of the sequence, it is mostly focusing on the nucleotides at the 3’ end, whereas the LSTM is such that is retains information from the 5’ end. However, the nucleotides at the 3’ end of the sgRNA sequence are known to carry the most influence over the activity of the sgRNA [13], and thus it is possible that in the LSTM nucleotides from the 5’ end are impacting the predicted activity more than they ought to. It would be worthwhile to run experiments with the sgRNA sequence reversed such that it runs 3’ to 5’.

The CNN LSTM+GC model actually showed slightly better performance than the other two in the activity range of 0 to 20%, then falling far behind for the rest of the activity deciles. The CNN GRU+GC and CNN BiLSTM+GC models performed quite similarly across all deciles, however, the CNN GRU+GC model slightly outperformed the CNN BiLSTM+GC model at higher activity deciles (70-100%) (Figure 7c).

### 3.6 Comparison to Other Models

Two other studies have published results of their models developed on this dataset exclusively: DeepHF and AttCRISPR. Shown in Table 4 is a comparison of our ChromeCRISPR models, including our top model, the hybrid CNN GRU+GC, with the LSTM models from DeepHF with and without bio-features, as well as the several attention-based models from AttCRISPR with and without bio-features. We found that our CNN GRU+GC ChromeCRISPR model outperformed both the base DeepHF and AttCRISPR models, as well as the ones with bio-features.

**Table 4:**
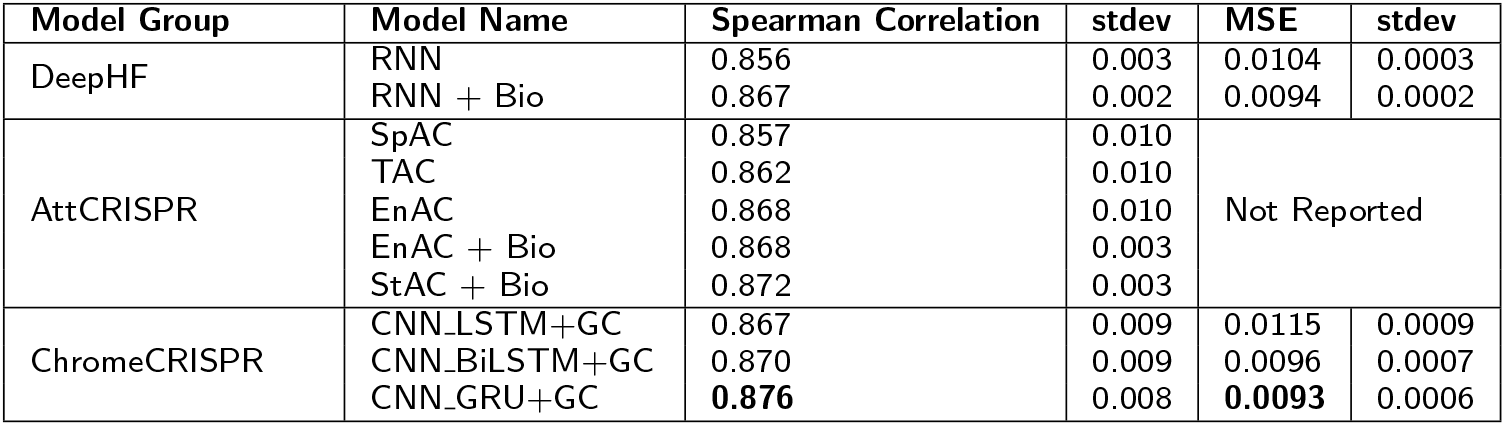
Comparison to other SotA Models.

### 4. Discussion

The strategic incorporation of GC content in the last fully connected layer of our model, just before the final prediction, represents a significant advancement in sgRNA activity prediction. This approach allows the network to initially learn complex representations of sgRNA sequences through multiple layers of abstraction. Integrating GC content at this later stage enables the model to combine learned sequence features with the biological relevance of GC content effectively. This nuanced method showcases the model’s ability to leverage both in-depth sequence understanding and critical biological indicators, like GC content, to enhance predictive accuracy.

Our research has been directed towards exploring the efficacy of hybrid models in sgRNA activity prediction, particularly through the integration of convolutional neural networks (CNNs) with recurrent neural network (RNN) architectures. This approach leverages the strengths of both CNNs, with their capability to extract spatial features from sequence data, and RNNs, notably GRU (Gated Recurrent Unit) layers, which excel in capturing temporal dependencies within sequences.

The motivation behind employing a hybrid CNN-GRU architecture stems from the complexity of sgRNA sequences, where both the sequence-specific features and the order of nucleotides play critical roles in determining sgRNA efficacy. CNN layers are adept at identifying relevant patterns across sgRNA sequences, while GRU layers contribute to understanding how these patterns influence activity outcomes based on their arrangement within the sequence.

In addition to our ChromeCRISPR hybrid models, we also experimented with a standalone Transformer architecture. However, this Transformer-based model did not outperform our deep RNN-based models. This suggests that the specific Transformer architecture we tested might not have been suitable for this task, or its known preference for longer sequences might not align well with the 21-mer sequences we have in this project.

### 4.1 Future Directions

#### 4.1.1 Adding Attention and Additional Bio-features

The attention mechanism has shown promise in various sequence-based prediction tasks, allowing models to focus on the most relevant parts of the input sequence [38, 39] and could be explored in our model and evaluated against current models such as [40]. CRISPR/Cas9 efficacy is influenced by numerous biological factors beyond the sgRNA sequence itself, such as the target gene’s expression level, chromatin accessibility, and the genomic context surrounding the target site [14,41]. To capture these influences, we intend to incorporate relevant biofeatures extracted from public genomic databases and the literature into our predictive models [42, 43].

#### 4.1.2 Embedding for Data Representation

The representation of sgRNA sequences is critical for accurately capturing the biological information essential for predicting CRISPR/Cas9 efficacy. Traditional one-hot encoding may not capture all relevant sequence characteristics. Therefore, we plan to investigate advanced embedding techniques, such as position-specific scoring matrices (PSSM) and word2vec models, for generating dense vector representations of sgRNA sequences [44]. These techniques are expected to provide a more nuanced representation of sgRNA sequences by encapsulating the identity, positional importance, and context of nucleotides within the sequence. The adoption of these advanced embeddings will be explored to uncover latent features correlating with on-target efficacy, with the aim of enhancing model performance and generalizability [45, 46].

#### 4.1.3 Generalizability of the Model

We would like to see how our model can generalize in two different cases: 1) To CRISPR/Cas9 data from other studies, and 2) To other Cas enzymes, e.g. eSpCas9 [47] and SpCas9-HF [48].

### 5. Conclusion

CRISPR/Cas is one of the leading gene editing methodologies and on-target prediction remains a challenging problem. This paper introduces *ChromeCRISPR*, a novel hybrid machine learning model that combines the strengths of CNNs with RNNs and leads to a high efficacy for CRISPR/Cas on-target predictions. The research presented here builds on our previous work. With the addition of deep models, hybridization and adding GC content we were able to outperform other state-of-the-art methods developed from the same dataset (i.e. DeepHF and AttCRISPR).

We achieved this through systematic experiments where we assessed the impact of the depth of the models and adding GC content as a feature. We analyzed how these factors affect the prediction of sgRNA activity values across sgRNAs with various levels of GC content and observed activity. From these analyses, we developed hybrid models that combined the feature extraction capabilities of the CNN with the powerful sequence processing of the RNNs. Our best model, a CNN-GRU hybrid with GC content was able to surpass previous models in terms of both the Spearman correlation and the mean squared error.

The significance of *ChromeCRISPR* lies in its potential to advance CRISPR/Cas-based therapies for genetic disorders and other gene editing experiments. Also, the study presented here presents insights into which machine learning techniques are best suited for this task. Designing highly effective sgRNAs that specifically target the defective DNA can lead to more precise and safe gene editing. This in turn could facilitate the development of treatments for previously incurable genetic diseases. Our study shows the power of hybrid models, as well as the positive impact of adding GC content as a feature. We hope that *ChromeCRISPR* will aid in the establishment of robust predictive models for CRISPR/Cas activity, which in turn could promise a transformative impact on human health and medical advancements.

## Author’s Contributions

The experimental design was a joint collaboration between AD and MF. AD implemented and ran the models, while MF analysed the results. KCW provided research supervision and guidance, and reviewed the data and analysis. All authors contributed to the writing and editing of the final manuscript.

## Acknowledgements

This project was supported in part by the Natural Sciences and Engineering Research Council of Canada (NSERC) under grant number RGPIN/04971-2020. We also acknowledge the Digital Research Alliance of Canada for providing computational resources to run our experiments.

